# Characterization of Telomeric Repeat-Containing RNA (TERRA) localization and protein interactions in Primordial Germ Cells of the mouse

**DOI:** 10.1101/362632

**Authors:** Miguel A. Brieño-Enríquez, Stefannie L. Moak, Anyul Abud-Flores, Paula E. Cohen

**Affiliations:** Department of Biomedical Sciences and Center for Reproductive Genomics, Cornell University, Ithaca, New York 14853, United States of America; Facultad de Medicina de la, Universidad Autónoma de San Luis Potosí, San Luis Potosí, México

**Keywords:** TERRA, Primordial Germ Cells, Long non-coding RNA, Telomeres, SFPQ, NONO

## Abstract

Telomeres are dynamic nucleoprotein structures capping the physical ends of linear eukaryotic chromosomes. They consist of telomeric DNA repeats (TTAGGG), the shelterin protein complex, and Telomeric Repeat-Containing RNA (TERRA). Proposed TERRA functions are wide-ranging and include telomere maintenance, telomerase inhibition, genomic stability, and alternative lengthening of telomere. However, the role of TERRA in primordial germ cells (PGCs), the embryonic precursors of germ cells, is unknown. Using RNA-fluorescence in situ hybridization (RNA-FISH) we identify TERRA in PGCs soon after these cells have migrated to, and become established in, the developing gonad. RNA-FISH showed the presence of TERRA transcripts in female PGCs at 11.5, 12.5 and 13.5 days post-coitum. In male PGCs, however, TERRA transcripts are observable from 12.5 dpc. Using qPCR we evaluated chromosome-specific TERRA expression, and demonstrated that TERRA levels vary with sex and gestational age, and that transcription of TERRA from specific chromosomes is sexually dimorphic. TERRA interacting proteins were evaluated using Identification of Direct RNA Interacting Proteins (iDRiP) which identified 48 in female and 26 in male protein interactors specifically within nuclear extracts from PGCs at 13.5 dpc. We validated two different proteins the splicing factor, proline- and glutamine-rich (SFPQ) in PGCs and Non-POU domain-containing octamer-binding protein (NONO) in somatic cells. Our results show that, TERRA interacting proteins are determined by sex in both PGCs and somatic cells. Taken together, our data indicate that TERRA expression and interactome during PGC development are regulated in a dynamic fashion that is dependent on gestational age and sex.

Research reported in this publication was supported in part by the Eunice Kennedy Shriver National Institute of Child Health & Human Development of the National Institutes of Health under Award Number K99HD090289 to M.A.B-E. A seed grant from the Cornell Center for Reproductive Genomics to M.A.B-E, using funds obtained as part of the NICHD National Centers for Translational Research in Reproduction and Infertility (NCTRI), award number P50HD076210 to P.E.C. and Empire State Stem Cell Fund through New York State Department of Health Contract # C30293GG. Imaging data was acquired through the Cornell University Biotechnology Resource Center, with NSF funding #1428922 for the shared Zeiss Elyra Microscope. NIH SIG 1S10 OD017992-01 grant support the Orbitrap Fusion mass spectrometer. The funders had no role in study design, data collection and analysis, decision to publish, or preparation of the manuscript.

## Introduction

Telomeres are dynamic nucleoprotein structures capping the physical ends of linear eukaryotic chromosomes. Telomeres protect the chromosome ends from degradation and erroneous recombination events and, as such, are essential for ensuring genome stability ^1^. They consist of DNA, proteins and RNA. Telomeric DNA consists of double strand DNA repeats (TTAGGG) which extend 9-15 kb in length in humans but can be as long as 100 kb in rodents ^2-4^. The protein component of the telomere is comprised of the shelterin complex, consisting of TRF1, TRF2, RAP1, TIN2, TPP1 and POT1 ^5^. The shelterin complex plays numerous roles in telomere stabilization, including suppression of the DNA damage response (DDR) and the regulation of telomerase activity ^6^. The Telomeric RNA component is TERRA (Telomeric repeat-containing RNA) a long non-coding RNA (lncRNA) ^3, 7-9^. Long non-coding RNAs are a class of RNAs defined by their size and by their lack of a translation product ^10^.

TERRA transcripts are comprised of UUAGGG repeats that are transcribed by RNA polymerase II, initiating from the subtelomeric regions of the telomeres and proceeding toward the chromosome ends ^5^. TERRA transcription in human cells is regulated by promoters localized at all subtelomeric regions, and by methylation on the CpG islands of these promoters ^11, 12^. By contrast, in mice only one promoter has been described on chromosome 18 and its regulation does not appear to dependent on methylation ^13^. TERRA expression levels and localization are cell cycle dependent, being high during the G1-S transition, peaking at early S phase and declining as cells transition to G2 and M phase ^11^. TERRA was initially described to be associated exclusively with the ends of linear eukaryotic DNA but recently studies have indicated that is also associates with other regions of the genome ^14^.

Several TERRA functions have been described, including telomere maintenance, telomerase inhibition, telomeric heterochromatin formation, genomic stability, an alternative lengthening mechanism for telomeres ^14-16^, and regulation of telomerase ^14, 17^. More recently, TERRA has been implicated in the protection of telomere stability, in which TERRA competes with ATRX to bind to DNA ^14^, and in the regulation of sex chromosome pairing during stem cell X chromosome inactivation, where TERRA creates a hub to guide X inactivation center homology searching ^18^.

TERRA localizes to the telomeres of mammalian germ cells (oocytes and spermatocytes) during all stages of meiotic prophase I, where it appears to be regulated in a sex-specific manner ^3, 19^. However, the function and timing of TERRA transcription remains unclear. Preliminary data suggest that TERRA transcription initiates in the primordial germ cell stage ^20^. Primordial Germ Cells (PGCs) are the embryonic precursors of the germ cell lineage that will form the oocytes and sperm required for sexual reproduction ^21, 22^. In mice, PGCs first become identifiable as a cluster of approximately 40 cells at the base of the allantois at around embryonic day 6.25 days post coitum (dpc). Starting around 7.5 dpc, PGCs begin their migration towards the presumptive gonad, colonizing the genital ridge by ~10.5 dpc ^21, 23^, and reaching their maximal numbers by ~13.5 dpc through continued migration and proliferation ^24^. In the female embryo, at 13.5 dpc PGCs then enter prophase I of meiosis and arrest at diplotene by around the time of birth. Meanwhile, male PGCs at this gestational age undergo mitotic arrest in the G_0_/G_1_ phase and stay quiescent for the remainder of the embryonic period, only initiating meiosis after birth ^21^.

In the current study, we have investigated the localization, transcription, and protein interactions of TERRA in male and female PGCs just after sex determination, and prior to the time that female germ cells enter meiosis and male germ cells become quiescent. Our results demonstrate key differences in the expression of TERRA and the localization of TERRA between male and female PGCs and across gestational age. Importantly, we also observe sex and age-dependent variations in the proteins with which TERRA interacts. Our results demonstrate that TERRA expression, localization and interactome in PGCs are sexually dimorphic and dependent on developmental age, suggesting that TERRA regulation may be an important component of the sex-specific development of the mammalian germ line.

## Materials and Methods

### Mouse Handling and Care

All mouse studies were conducted with the prior approval of the Cornell Institutional Animal Care and Use Committee. 6 to 8 week old wild-type C57Bl/6J mice were mated and checked for vaginal plugs the next morning. Female mice with plugs were moved into a separate cage, and were considered to be at 0.5 days post-coitum (dpc). At 11.5, 12.5 and 13.5 dpc, the mothers were sacrificed to recover gonads from fetuses for the isolation of PGCs.

### Extraction of Primordial Germ Cells

PGCs were purified by magnetic cell sorting of gonads from 11.5, 12.5 and 13.5 dpc male and female embryos following a published protocol with some modifications ^25, 26^. Briefly, the gonads from 11.5, 12.5 and 13.5 dpc embryos were dissected under a stereomicroscopy in EmbryoMax M2 Medium (Merck-Millipore). Gonads at 12.5 dpc and 13.5 dpc were classified according to the anatomical characteristics. For 11.5 dpc gonads dissected and sexed by PCR following the method described by McClive, P.J. & Sinclair ^27^. Testis and ovaries were disaggregated in 0.25% trypsin-EDTA containing 20 µg/ml DNase (Sigma-Aldrich, St. Louis, MO. USA) at RT. The enzymatic reaction was stopped by adding M2 medium with 10% FCS (Gibco, Thermo Fisher, Waltham, MA. USA). The cells were centrifuged at 3220 rcf for 2 min and washed twice with 500 µl of M2 medium containing 10% FCS and 20 µg/ml DNase. The cell pellet was resuspended in 400 µl of M2 medium, mixed with 30 µl anti-SSEA-1 (CD15) MicroBeads (Miltenyi Biotech, Bergisch Gladbach, Germany) and incubated for 45 min at 4°C. PGCs were isolated from somatic cells with a miniMACs column following the manufacturer’s instructions and counted (Miltenyi Biotech, Bergisch Gladbach. Germany). The purity of PGCs was verified by, counting cells that stained positive for alkaline phosphatase with the naphtol AS-MX/FAST-RED procedure (Sigma-Aldrich, St. Louis, MO. USA). In all cases the purity of the PGCs was evaluated from cell counts in 4 different fields of the microscope until a total number of 100 cells were recorded. The purity of PGCs varies according to gestational age: at 11.5 dpc, cell suspensions showed a purity of 80–85%, at 12.5 dpc the purity of PGCs ranged from 85 to 89%, and at 13.5 dpc PGC there was a recorded 90–95% purity. Typical yield of PGCs per embryo varies by gestational age. At 11.5 dpc the total isolated PGCs was approximately 800, at 12.5 dpc there were approximately 4000 PGCs isolated per embryo and approximately 10000 PGCs per embryo at 13.5 dpc. The somatic cells that were separated in the MS column were also collected for analysis.

### Primordial germ cell (PGCs) and somatic cell spreading (Soma)

After extraction, 20 µl of the cell suspension was placed on poly-L-lysine slides that were previously cleaned with RNaseZap (Thermo Fisher, Waltham, MA USA). Cells were permeabilized by incubation for 10 minutes in CSK buffer (100 mM NaCl, 300 mM sucrose, 3 mM MgCl2, 10 mM PIPES (Sigma-Aldrich, St. Louis, MO. USA), 0.5% Triton X-100 (Fisher Scientific, Pittsburgh, PA. USA) and 10 mM ribonucleoside-vanadyl complex (New England Biolabs, Ipswich, Massachusetts. USA)) ^28^. Cells were fixed in 4% paraformaldehyde (Electron Microscopy Sciences, Hatfield, PA. USA) for 10 min and then washed in 70% ethanol. Slides were stored at −80°C until use.

### Immunofluorescence

Immunofluorescence (IF) was performed as previously described with modifications ^28^. Slides were blocked 10 min with PTBG (1x PBS, 0.1% Tween-20, 0.2% BSA, 0.2% gelatin) and then incubated overnight at 4°C with mouse anti*-Tert* (dilution 1:100 from Rockland antibodies #600-401-252S), anti-NONO (dilution 1:100 from Proteintech #11058), anti-SFPQ (dilution 1:100 from ProteinTech #15585) and anti-VASA (DDX4) (dilution 1:200 from Abcam #ab13840) in PTBG. Slides were washed three times for 5 minutes in PBST (0.1% Tween-20 in 1x PBS) before incubation at 37°C for 40 minutes with the following secondary antibody: Alexa Fluor 488-AffiniPure F(ab’)2 Fragment goat anti-mouse IgG (H+L), Alexa Fluor 488-AffiniPure F(ab’)2 Fragment goat anti-rabibit IgG (H+L) and Alexa Fluor 594-AffiniPure F(ab’)2 Fragment goat anti-rabbit IgG (H+L), all of them from Jackson Immunoresearch (West Grove, PA. USA). Secondary antibodies were incubated at concentrations of 1:1000 in PTBG at 37°C for 60 minutes. Slides were washed three times in PBST, fixed 10 min in 4% paraformaldehyde in 1x PBS (pH 7), and rinsed with 1x PBS. At the end of the IF procedure, RNA-FISH was performed.

### RNA-Fluorescence In Situ Hybridization

TERRA focus detection was performed by RNA-FISH ^28, 29^. RNA-fluorescence in situ hybridization (RNA-FISH) was performed immediately after IF. Briefly, after dehydration through a ice cold graded ethanol series (70%, 80%, 90% and 100%, during 5min each), cells were hybridized overnight at 37°C with a 25 nM (CCCTAA)^3^ oligonucleotide probe conjugated with Cy3 (Integrated DNA Technologies IDT, Coralville, IO. USA) in hybridization buffer (10% of 20x SSC (Sigma-Aldrich, St. Louis, MO. USA), 20% 10 mg/ml BSA (Sigma-Aldrich, St. Louis, MO. USA), 20% of 50% dextran sulfate (Fisher Scientific, Pittsburgh, PA. USA), and 5% deionized formamide (Fisher Scientific, Pittsburgh, PA. USA). Next, slides were washed with 50% formamide/1xSSC during 5 min followed by two washes of 2xSSC at 39°C. Finally, cell nuclei were counterstained with 4,6-diamidino-2-phenylindole (DAPI) diluted in Vectashield (Vector Laboratories, Burlingame, CA). For each sample, a negative control was included, consisting of slides treated 10 min with RNAse A (Sigma-Aldrich, St. Louis, MO. USA) at a concentration of 100 µg/ml prior to RNA-FISH performance. Foci were counted as signals that appeared as discrete dots or signals within the cell. At least 75 cells were counted per each PGC pool and 50 for each somatic cells pool.

### Immunohistochemistry

Immunohistochemistry was performed on paraffin-embedded sections from 13.5 dpc ovaries and testis. Slides were deparaffinized and rehydrated with 3 washes of Safeclear (Fisher Scientific, Pittsburgh, PA. USA) for 5 minutes each, followed by 3 washes of each concentration in a graded series of ethanol (100%, 95%, 80%, 70%). After rinsing the slides 2 times for 5 minutes in distilled water, the slides were incubated in sodium citrate pH 6.0 during 40 min at 95 °C. Permeabilization was performed in 0.2% of Triton-X 100 in PBS for 30 min. Section were blocked for 4 h in blocking solution (2.52 mg/ml glycine,10% goat serum, 3% BSA in PBS-T) and then incubated with primary antibody overnight at RT. After 2 washes with PBST, the slides were incubated with secondary antibodies for 2 h at RT. The slides where rinsed in PBST, cell nuclei were counterstained with 4,6-diamidino-2-phenylindole (DAPI) and mounted in Vectashield (Vector Laboratories, Burlingame, CA).

### Imaging

Imaging of PGCs and somatic cells was performed using ELYRA 3D-Structured Illumination Super resolution Microscopy (3D-SIM) from Carl Zeiss with ZEN Black software (Carl Zeiss AG, Oberkochen. Germany). Images are show as maximum intensity projections of z-stack images-3D-SIM. To reconstruct high-resolution images, raw images were computationally processed by ZEN Black. Channel alignment was used to correct for chromatic shift. The brightness and contrast of images were adjusted using ImageJ (National Institutes of Health, USA). Image acquisition of the tissue sections was performed using a Zeiss Imager Z1 microscope under 20X, 40X or 63X magnifying objectives, at RT. Images were processed using ZEN 2 (Carl Zeiss AG, Oberkochen. Germany).

### RNA isolation and reverse transcription

Total RNA from each gestational age (11.5, 12.5 and 13.5 dpc for both male and female gonads) was extracted using TRIzol (Invitrogen Co, Carlsbad, CA, USA). Total RNA was re-suspended in 40 µl of RNase free water. RNA concentration was then determined spectrophotometrically using a NanoDrop 2000 (Thermo Fisher, Waltham, MA. USA). 1.5µg of total RNA was reverse transcribed using and Superscript III First-Strand Synthesis System (Invitrogen Co, Carlsbad, CA, USA) using TERRA reverse primer or 1.65 μM random hexamers/1.25 μM Oligo(dt) for the housekeeping gene (Supplemental Table 1). cDNA was kept at −20°C until used in qPCR.

### qPCR analysis

Real-time PCR amplification and analysis was performed following the protocol previously described by Feretzaki and Lingner ^30^. Primers were designed to amplify the subtelomeric region of different chromosomes using the sequences previously published by Lopez de Silanes et al., 2014 ^13^ (Supplemental Table 1) (Subtelomeric regions with active transcription of functional genes were not included). Gene expression was normalized to succinate dehydrogenase complex flavoprotein subunit A (SDHA) expression ^31^. Each reaction mixture consisted of, 1 µl of cDNA, 0.5 µl of forward primer (0.2 µM), 0.5 µl of reverse primer (0.2 µM), 10 µl of Roche FastStart SYBR Green Master (Sigma-Aldrich, St. Louis, MO. USA) and 7 µl of nuclease-free water. qRT-PCR amplification reaction was performed with specific primers (Supplemental Table 1). PCR conditions were the same as those used by Feretzaki and Lingner ^30^: 30 s at 95°C, followed by 40 cycles at 95°C for 1 s and 60°C for 60 s. After PCR, melting curve analyses were performed to verify specificity and identity of the PCR products. All data were analyzed with the CFX-manager Bio-Rad (Bio-Rad Laboratories). All analyzed genes were performed in triplicate for each one of the 3 biological samples of the three different developmental ages (11.5, 12.5 and 13.5 dpc) and both sexes (female and male). qPCR data for TERRA quantification are analyzed using the relative quantification method ^32^. This method feeds the Ct values obtained from the qPCR experiment into a series of subtractions to calculate the relative gene expression of the gene of interest (TERRA) normalized against a reference gene (SDHA) in different conditions as was described previously by ^14, 18, 30^.

### Identification of Direct RNA Interacting Proteins (iDRiP)

Proteins that directly interact with TERRA were identified through method called iDRiP, following a published protocol with some adaptations ^33^. The original iDRiP method utilized large numbers of cultured somatic cells (around 30 million). In the current study, we adapted the conditions to reduce the input of cells around 10-fold and to use primary PGCs from 13.5 dpc male and female gonads, along with gonadal somatic cells as controls. After cell isolation, cells were rinsed with cold PBS 3 times and the plated in a petri dish for 30 min at 37°C 5% CO_2_. Excess PBS was removed and the cells were irradiated with UV light at 200 mJ/cm^2^ energy (Spectrolinker XL, UV crosslinker. Spectronics Corporation. Westbury, NY. USA). The cells were transferred to an eppendorf tube and spun down at 1258 rcf for 5 min at 4°C. The pellet was re-suspended in 300 µl cold cytoskeleton buffer (CSKT) (0.5% Triton X-100, 0.5% PIPES, 100 mM NaCl, 3 mM MgCl2, 0.3 M sucrose and 1 mM PMSF) (all from Fisher Scientific, Pittsburgh, PA. USA) with protease inhibitor (1x Roche Protease Inhibitor Cocktail Tablets) and incubated for 10 min at 4°C on a rocker. Cells were spun down again at 453 rcf at 4°C for 5 minutes, supernatant was removed and re-suspended in nuclear isolation buffer (10 mM Tris pH 7.5, 10 mM KCl, 0.5% Nonidet-P 40, 1x protease inhibitors, 1 mM PMSF), before spinning again at 453 rcf at 4°C for 10 min. Supernatant was removed and the cell pellets were flash frozen in liquid nitrogen and stored at −80°C until use. Cells were pooled from number of ≅300 female and ≅300 male embryos and thawed at 37°C. 500 µl Turbo DNaseI buffer, 50 µl Turbo DNaseI enzyme (2U/µl), 10 µl superaseIN (Thermo Fisher, Waltham, MA USA) and 5 ul of 50x protease inhibitor was added to the cell suspension. Samples were incubated at 37°C for 45 min on a rocker. The nuclear lysates were further solubilized by adding 1% sodium lauryl sarcosine, 1x protease inhibitor, 0.3 M lithium chloride, 25 mM EDTA and 25 mM EGTA to final concentrations (all of them from Sigma-Aldrich, St. Louis, MO. USA). Samples were mixed well and incubated again at 37°C for 15 min. As a positive control, the highly expressed RNA U6 was used, and RNAse A treated samples as a negative control. U6 and TERRA-specific biotinylated probes (Integrated DNA Technologies IDT, Coralville, IO. USA), were conjugated to streptavidin beads (MyOne streptavidin C1 Dyna beads, Invitrogen) for a 30 min incubation period at RT. The conjugated beads were mixed with the lysates and incubated at 55°C for one hour before overnight incubation at 37°C in a hybridization chamber. After incubation, the beads were washed three times in wash buffer (10 mM Tris, pH 7.5, 0.3 M LiCl, 1% LDS, 0.5% Nonidet-P 40, 1x protease inhibitor) at RT then treated with DNase I digestion buffer, Turbo DNase I, 0.3 M LiCl, protease inhibitors, and SuperaseIn (Thermo Fisher, Waltham, MA USA) at 37°C for 20 min. Beads were re-suspended and washed two more times in the wash buffer. For mass spectrometry analysis, proteins were eluted in Elution Buffer (10 mM Tris, pH 7.5, 1 mM EDTA) at 70°C for 4 min.

### Nano-scale reverse phase chromatography and tandem MS (nanoLC-MS/MS)

Mass spectrometry of iDRiP-derived proteins was performed in the Cornell University Proteomics and Mass Spectrometry facility. The nanoLC-MS/MS analysis was carried out using an Orbitrap Fusion (Thermo-Fisher Scientific, San Jose, CA) mass spectrometer equipped with a nanospray Flex Ion Source using high energy collision dissociation (HCD) and coupled with the UltiMate3000 RSLCnano (Dionex, Sunnyvale, CA). Each reconstituted samples for both PGC and SOMA (18 ul) was injected onto a PepMap C-18 RP nano trap column (3 µm, 100 µm × 20 mm, Dionex) with nanoViper Fittings at 20 μL/min flow rate for on-line desalting and then separated on a PepMap C-18 RP nano column (3 µm, 75 µm x 25 cm), and eluted in a 120 min gradient of 5% to 35% acetonitrile (ACN) in 0.1% formic acid at 300 nL/min. The instrument was operated in data-dependent acquisition (DDA) mode using FT mass analyzer for one survey MS scan for selecting precursor ions followed by 3 second “Top Speed” data-dependent HCD-MS/MS scans in Orbitrap analyzer for precursor peptides with 2-7 charged ions above a threshold ion count of 10,000 with normalized collision energy of 38.5%. For label-free protein analysis, one MS survey scan was followed by 3 second “Top Speed” data-dependent CID ion trap MS/MS scans with normalized collision energy of 30%. Dynamic exclusion parameters were set at 1 within 45s exclusion duration with ±10 *ppm* exclusion mass width. Two samples from each group PGC and SOMA were analyzed in Orbitrap in the order of female followed by male samples for data acquisition. All data are acquired under Xcalibur 3.0 operation software and Orbitrap Fusion Tune 2.0 (Thermo-Fisher Scientific).

### NanoLC-MS/MS data processing, protein identification and data analysis

All MS and MS/MS raw spectra from each experiment were processed and searched using the Sequest HT search engine within the Proteome Discoverer 2.2 (PD2.2, Thermo). The default search settings used for relative protein quantitation and protein identification in PD2.2 searching software were: two mis-cleavage for full trypsin with fixed carbamidomethyl modification of cysteine and oxidation of methionine and demaidation of asparagine and glutamine and acetylation on N-terminal of protein were used as variable modifications. Identified peptides were filtered for maximum 1% false discovery rate (FDR) using the Percolator algorithm in PD 2.2. The relative label free quantification method within Proteome Discoverer 2.2 software was used to calculate the protein abundances. The intensity values of peptides, which were summed from the intensities values of the number of peptide spectrum matches (PSMs), were summed to represent the abundance of the proteins. For relative ratio between the two groups, here PGC female/male and Soma female/male, no normalization on total peptide amount for each sample was applied. Protein ratios are calculated based on pairwise ratio, where the median of all possible pairwise ratios calculated between replicates of all connected peptides.

### Statistical Analysis

Statistical analyses were performed using GraphPad Prism version 6.00 for Macintosh (GraphPad Software, San Diego California USA, www.graphpad.com). Specific analyses are described within the text and the corresponding figures. Alpha value was established at 0.05.

## Results

### Developmental changes in TERRA localization are different in male and female PGCs

Using RNA-FISH and 3D-SIM microscopy, we evaluated the presence of TERRA at three developmental stages (11.5 12.5 and 13.5 dpc) in both sexes. The selection of these stages was based on the timing of sex determination in the mouse, the methylation status of PGCs, and the entrance into meiosis of female PGCs after 13.5 dpc ^21^. Quantitation of TERRA focus numbers was performed in both male and female SSEA-1 positive PGCs as well as in SSEA-1 negative cells (somatic cells). As a negative control, cells treated with RNAse A were used in which TERRA RNA should be completely degraded. A total number of 1352 PGCs and 973 somatic cells were analyzed. Female and male PGCs, and somatic cells were obtained from at least 3 different pools at the different gestational ages. Pools consisted of PGCs from between 20 and 40 gonad pairs (1 pair per embryo).

Analysis of female PGCs and somatic cells showed the presence of discrete foci of TERRA (Fig. 1A and 1B). The mean focus number observed in 11.5 dpc female PGCs was 0.76 ± 0.37 per cell, rising to 3.88 ± 0.73 at 12.5 dpc, and 8.76 ± 4.03 at 13.5 dpc. Statistical analysis revealed significant differences among the three gestational ages (p=0.001 ANOVA; Fig. 1B). Meanwhile, TERRA focus counts from female gonadal somatic cells at the same gestational ages revealed no statistical differences from 11.5 dpc (19.76 ± 1.143), 12.5 dpc (19.86 ± 1.52), and 13.5 dpc (19.78 ± 1.44; Fig. 1C and 1D). The number of TERRA foci in somatic cells was considerably higher than that observed in the neighboring germ cells at each gestational age. These results indicate that TERRA focus numbers alter with gestational in female PGCs but not in neighboring somatic cells.

**Figure 1.**
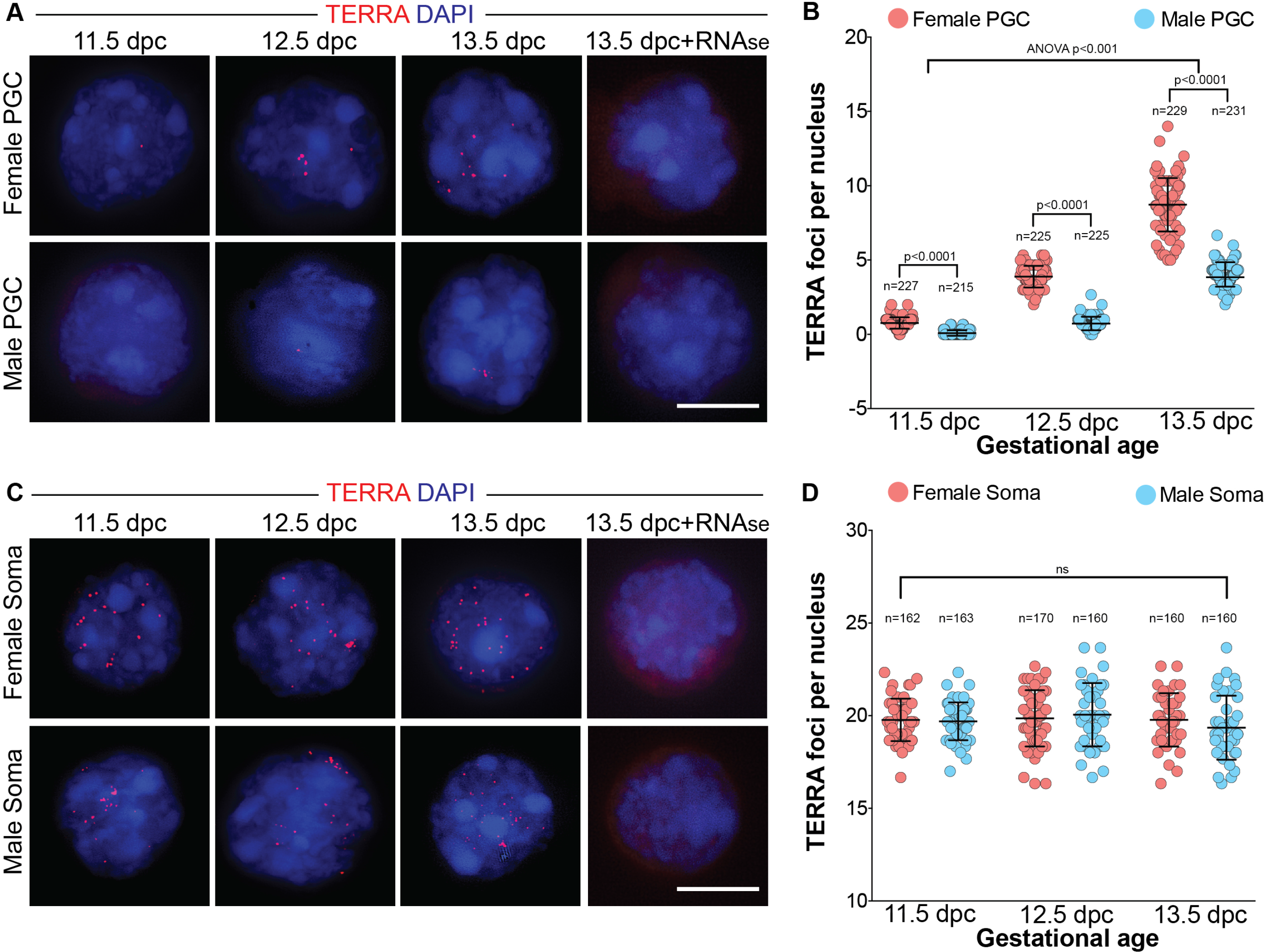
TERRA presence in mouse PGCs differs by sex and developmental age. A) Super resolution imaging microscopy of TERRA RNA-FISH (red) on female and male primordial germ cells at different stages of development (11.5, 12.5 and 13.5 dpc). Control negative correspond to 13.5 dpc female and male PGCs treated with RNAse before the RNA-FISH procedure. Scale bar 10 µm; B) Quantitation of TERRA foci numbers on female and male PGCs at different stages of development(11.5, 12.5 and 13.5 dpc). Statistical analysis was performed using ANOVA followed by Kruskal-Wallis multiple comparison analysis; C) Super resolution imaging microscopy of TERRA RNA-FISH (red) on female and male somatic cells at different stages of development (11.5, 12.5 and 13.5 dpc). Negative controls used were 13.5 dpc female and male somatic cell treated with RNAse before the RNA-FISH procedure. Scale bar 10 µm; D) Quantitation of TERRA foci numbers on female and male somatic cells at different stages of development (11.5, 12.5 and 13.5 dpc). Statistical analysis was performed using ANOVA followed by Kruskal-Wallis multiple comparison analysis. p value was set at 0.05.

In male PGCs, the dynamics of TERRA focus accumulation were very different to that seen in females. We were not able to detect TERRA signal in male 11.5 dpc PGCs. Instead, the earliest detection TERRA in male PGCs was at 12.5 dpc where we observed cells with either zero or one TERRA focus (0.73 ± 0.45 foci/cell). A statistically significant increase in the TERRA foci was observed in male 13.5 dpc PGCs, where the mean focus number rose to 4.03 ± .82 (p=0.001; Fig. 1A). Adjacent somatic cells of the male gonad showed TERRA focus numbers that were indistinguishable at all ages from that of female gonads, with no statistically significant differences found between sex or gestational age (11.5 dpc: 19.66 ± 1.02 foci/cell, 12.5 dpc: 20.05 ± 1.71 foci/cell, and 13.5 dpc: 19.35 ± 1.76 foci/per cell; Fig. 1D). These results indicate that in male PGCs, like in female PGCs, the gestational age is a key factor in TERRA localization in PGCs but not in somatic cells.

Comparison of TERRA foci numbers in both female PGCs and male PGCs reveled a sex bias. Compared to male PGCs, female PGCs showed significantly more TERRA foci at all the stages of development (p=0.001; Fig. 1B), while no differences were observed at any gestational age between male and female somatic cells (Fig. 1D).

### Differential transcription of TERRA in male and female PGCs

In human cell lines, transcription of TERRA is regulated by chromosome specific promoters that are repressed by CpG methylation ^34, 35^. However, previous reports in mouse cell lines showed that almost all the TERRA transcripts are transcribed from the subtelomeric region of chromosome 18, with some minor contribution from chromosome 9 ^13^. However, nothing is known about TERRA transcription during this unique period of PGC development in which distinct changes in DNA methylation are occurring. Thus, we hypothesized that the loss of methylation that is a unique feature of PGCs may result in increased TERRA transcription, resulting in the increased localization of TERRA that we observed through this gestational time span. We performed qPCR in isolated female and male PGCs at 11.5, 12.5 and 13.5 dpc using the subtelomeric sequences of each chromosome previously published by Lopez de Silanes et al., 2014 ^13^ (Supplemental Table 1). As expected from the localization of TERRA, our results showed higher levels of TERRA expression in female PGCs compared to male PGCs at each gestational age (Fig. 2A-2O). Furthermore, and in contrast to previous results, we observed that TERRA is transcribed from multiple telomeres in a gestational and sex-dependent manner, though not all telomeres were found to be transcriptionally active (only those with any expression of TERRA are shown in Figure 2).

**Figure 2.**
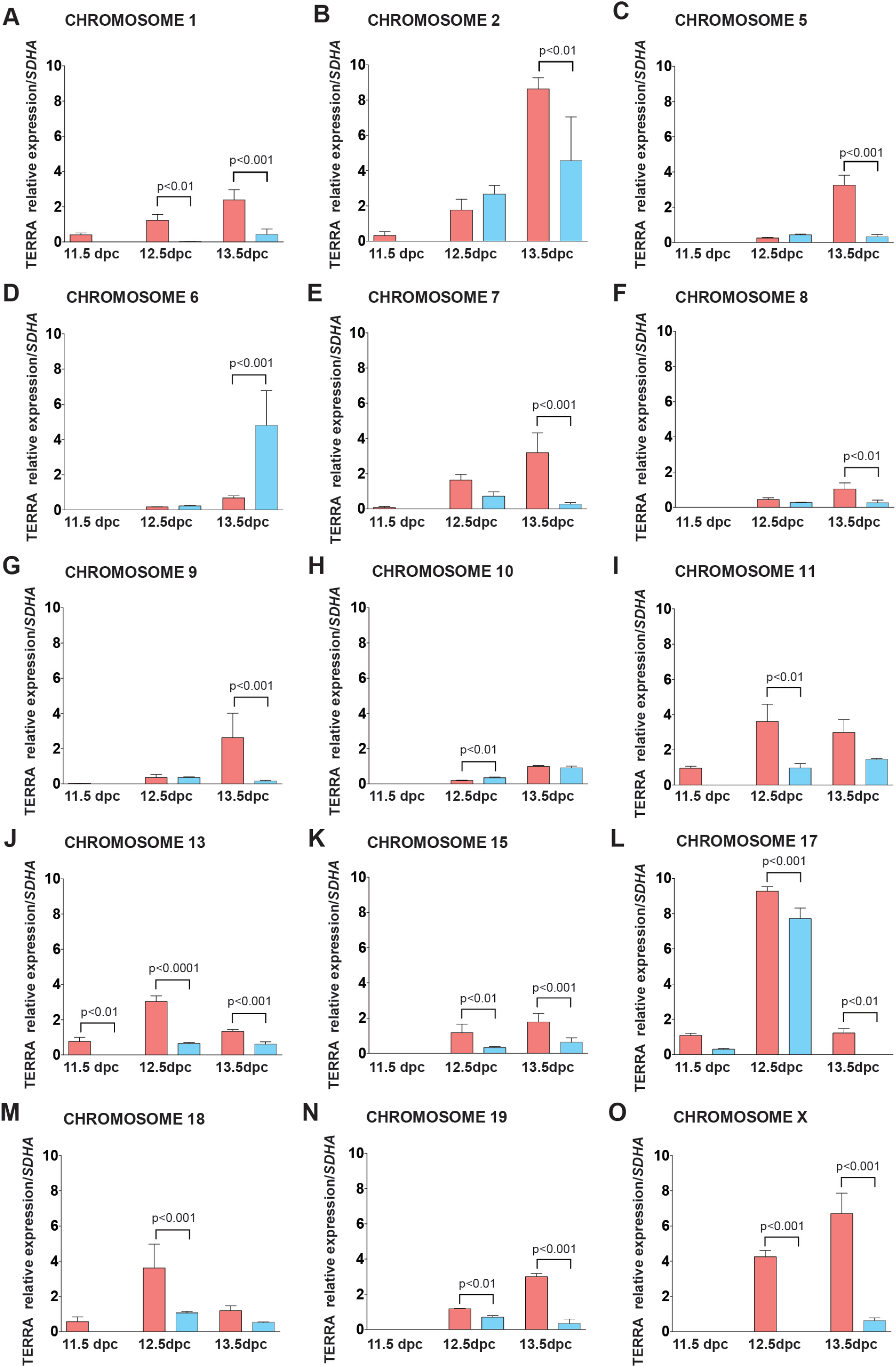
TERRA expression is different depending on gestational age and sex. A to O, qPCR analysis of the expression of TERRA from subtelomeric regions of chromosome 1, 2, 5, 6, 7, 8, 9, 10, 11, 13, 15, 17, 18, 19 and X. The results are expressed as relative TERRA expression normalized to *Sdha* at the different developmental ages (11.5, 12.5 and 13.5 dpc). Columns in red represent female PGCs and blue represent male PGCs Statistical analysis was performed using one-way ANOVA with multiple comparison. p value was set at 0.05

Female PGCs showed TERRA expression at 11.5 dpc from 8 different chromosome subtelomeric regions (Chromosomes 1, 2, 7,9, 11,13, 17 and 18; Fig. 2A, 2B, 2E, 2G, 2I, 2J, 2L and 2M, respectively), while in male PGCs at the same age, TERRA expression was confined to the chromosome 17 subtelomeric region (Figure 2L). At 12.5 dpc, we detected increased expression from chromosomes 5, 6, 8, 10, 15, 19 in both female and male PGCs (Fig. 2C, 2D, 2F, 2H, 2K and 2N, respectively). The exception was the subtelomeric region of chromosome X, from which transcription of TERRA was only detected for female PGCs (Fig. 2O). Transcription of TERRA from the single X chromosome of male PGCs only became evident at 13.5 dpc, but at a very low level compared to that of female PGCs at this gestational age (Fig, 2O). Most TERRA transcription in 13.5 dpc male PGCs arose from chromosomes 2 and 6, and only in the case of the latter was there higher transcription in male PGCs than female PGCs (Fig. 2B, 2D). Indeed, at all gestational ages, we observed higher transcription of TERRA at each telomere in female PGCs, except for the subtelomeric region of chromosomes 2, 6 and 10. These results indicate that the transcription of TERRA is differentially regulated in the male and female germ line, and that transcription of TERRA in male PGCs is developmentally delayed compared to that in female PGCs. Moreover, our results demonstrate that TERRA is transcribed from multiple subtelomeric regions in mouse PGCs.

### TERRA and the catalytic subunit of the enzyme telomerase (TERT) colocalization and expression are regulated by gestational age and sex

There is a lot of controversy regarding the role of TERRA in the regulation of Telomerase (TERT). The most common idea suggests that TERRA expression down-regulates or inhibits the catalytic subunit of the TERT ^17, 36^. Previous reports have indicated that there is decay in the *Tert* expression with age in male germ cells, including PGCs ^37, 38^. Confounding this, however, are the suggestions that high levels of telomerase are required to maintain spermatogonia in their undifferentiated state ^39^. Using SIM microscopy we evaluated the colocalization of TERT and TERRA at 13.5 dpc, and plotted the percent colocalization in PGCs from both sexes (percentages were obtained from the number of TERRA-TERT foci divided by the total TERRA foci, and multiplied by 100) (Fig. 3A and 3B). In female PGCs, 63.5% of the TERRA foci colocalized with TERT but only the 36.1% in 13.5 male PGCs (Fig. 3C; p=0.0001). We evaluated the expression of *Tert* using qPCR, and showed a statistically significant decrease of *Tert* expression 11.5 dpc to 13.5 dpc in both male and female PGCs (Fig. 3D; p=0.001). Consistently, however, *Tert* expression was significantly higher in male PGCs than in age-matched female PGCs (Fig. 3D). Thus, the increasing TERRA focus count with gestational age, and the relatively increased number of TERRA foci/cell in female germ cells compared to male germ cells are both inversely correlated with *Tert* expression, which declines with gestational age and which is higher in male PGCs.

**Figure 3.**
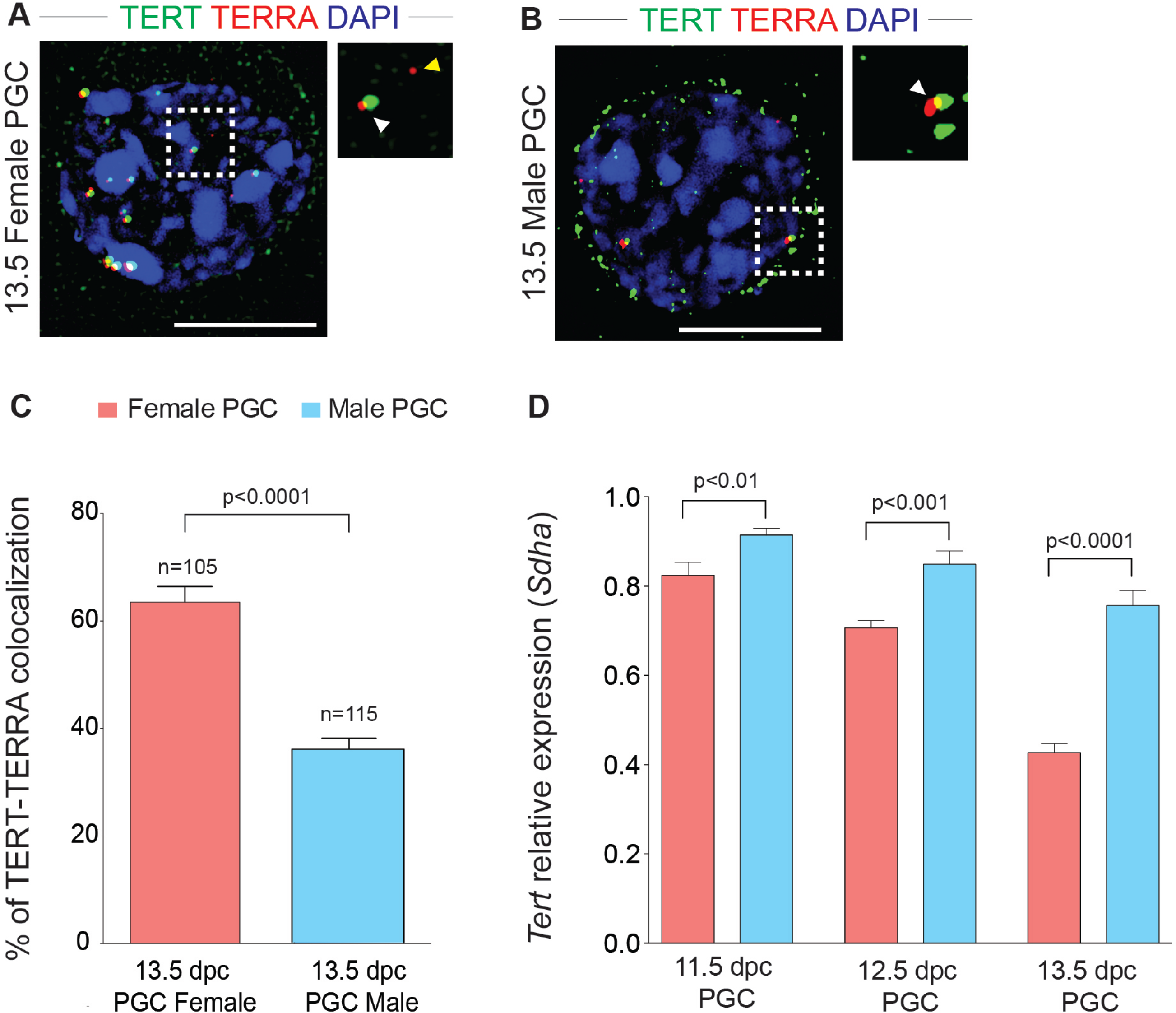
TERRA and the catalytic subunit of the enzyme telomerase (TERT) colocalization and expression are regulated by gestational age and sex. A) Super resolution imaging microscopy of TERT (green) immunofluorescence followed by TERRA RNA-FISH (red) on female 13.5 dpc PGCs. Scale bar 10 µm; in the inset the white arrowhead shows TERRA-TERT interaction, yellow arrow head shows only TERRA foci; B) Super resolution imaging microscopy of TERT (green) immunofluorescence followed by TERRA RNA-FISH (red) on male 13.5 dpc primordial germ cells. Scale bar 10 µm; in the inset the white arrowhead shows TERRA-TERT interaction, yellow arrow head shows only TERRA foci;C) Analysis of the percentage of TERRA-TERT colocalization on female and male PGCs. Percentages were obtained from the number of TERRA-TERT foci divided by the total TERRA foci, and multiplied by 100. Statistical analysis was performed with unpaired t-test. p value was set at 0.05; D) qPCR analysis of the expression of *Tert* at different stages of development (11.5, 12.5 and 13.5 dpc). The results are expressed as the *Tert* relative expression normalized by *Sdha*. Red columns indicated the analysis performed in female PGCs and blue columns indicate the analysis performed in male PGCs. Statistical analysis was performed using one-way ANOVA with multiple comparisons. p value was set at 0.05

### The TERRA interactome is sexually dimorphic

To understand TERRA function, it is important to identify key TERRA interacting proteins in the cell type of interest. Given that we obtain so few PGCs at specific developmental ages, we focused on only one gestational age in which to investigate the TERRA interactome in PGCs and somatics cells: 13.5 dpc. We performed iDRiP (Identification of Direct RNA Interacting Proteins), using three million PGCs from both male and female gonads, along with comparable neighboring somatic cells. A total of 48 proteins were identified in female PGCs and 26 in male PGCs, of which 32 (55.2%) were unique to female PGCs and 10 (17.2%) were specific for male PGCs (Fig. 4A and 4B). The remaining 16 (27.6%) TERRA-associated proteins were shared between female and male PGCs (Fig. 4A). Using the relative label free quantification method within Proteome Discoverer 2.2 software, we calculated the protein abundances. The intensity values of peptides, which were summed from the intensities values of the number of peptide spectrum matches (PSMs), were summed to represent the abundance of the proteins. The results of these relative label free quantitations showed different relative levels of protein between female and male PGCs (Fig. 4B) (Supplemental Table 2).

**Figure 4.**
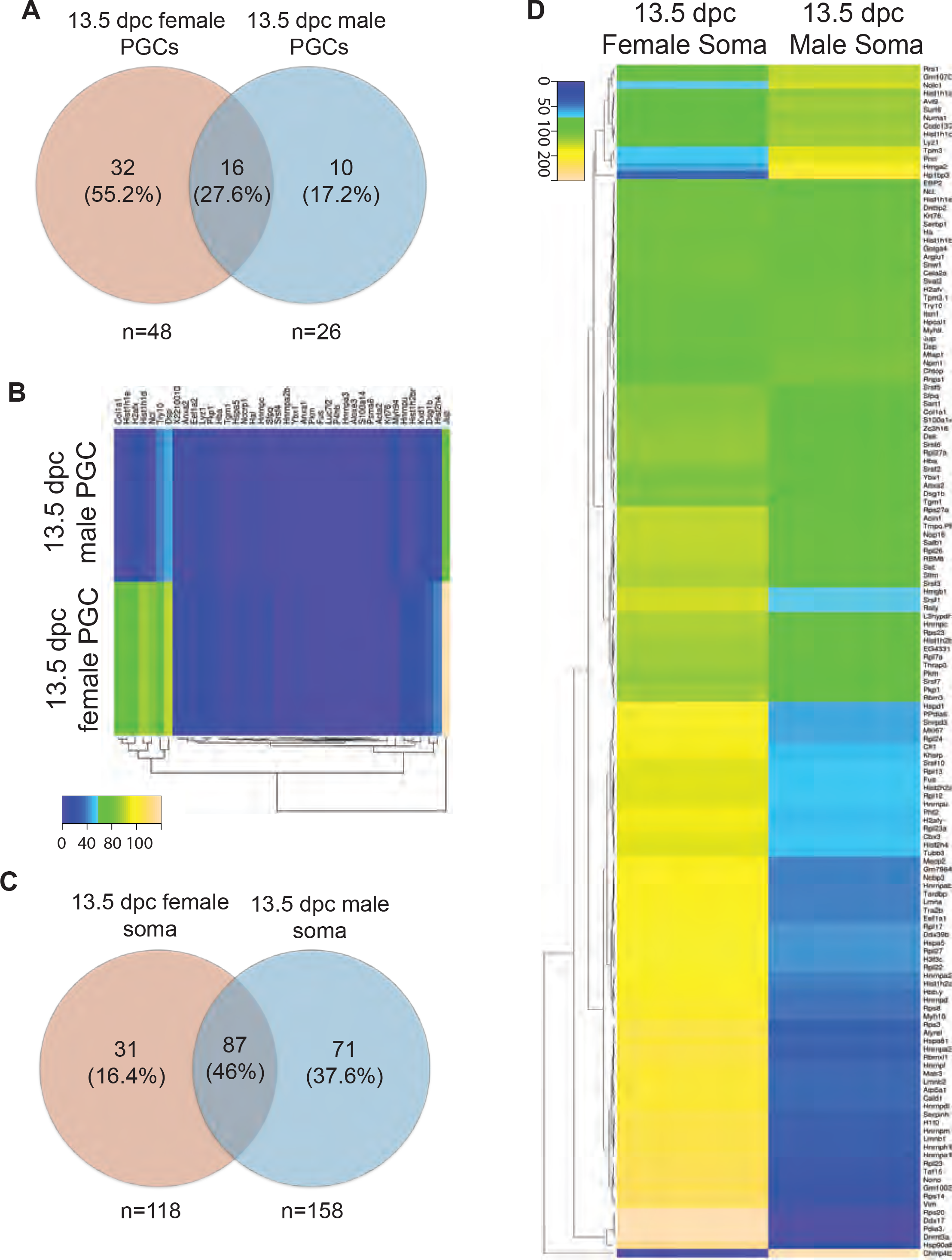
The TERRA interactome of the gonad is sexually dimorphic. A) Analysis of the male and female PGCs protein distribution obtained from iDRiP; B) Heat map analysis of the relative protein concentration in PGCs obtained by iDRiP-mass spec; The relative label free quantification method within Proteome Discoverer 2.2 software was used to calculate the protein abundances. C) Analysis of the male and female somatic protein distribution obtained from iDRiP; D) Heat map analysis of the relative protein concentration in somatic obtained by iDRiP-mass spec. The relative label free quantification method within Proteome Discoverer 2.2 software was used to calculate the protein abundances.

In somatic cells, we observe far higher numbers of TERRA interacting proteins, but with greater number of proteins in male somatic cells than in female somatic cells (Fig. 4C). 118 proteins were obtained from female somatic cells and 158 were obtained from male somatic cells (Fig. 4C), with 31 (16.4%) and 71(37.6%) proteins, respectively, being unique to one sex. Overall, female and male somatic cells shared 87 (46%) TERRA interacting proteins (Fig. 4D). Similar to PGCs, we used the relative label free quantification method to compare the relative abundance of proteins in both female and male somatic cells. Our results showed a different distribution of the relative protein abundance in female gonads compared to male gonads (Fig. 4D, and Supplemental Table 3).

### SFPQ interacts with TERRA in PGCs

To validate the interactions of TERRA we used two different approaches. First, we immunolocalized the protein of interest on 13.5 dpc ovaries and testis, and secondly, we co-localized TERRA and the protein on isolated cells. Based on our iDRIP-identified interacting proteins, we decided to analyze proteins with the most extreme ratios between male and female. After antibody testing and standardization, we selected two proteins for validation one for PGCs and another for somatic cells. The splicing factor, proline- and glutamine-rich (SFPQ) showed one of the lowest male/female ratios (0.01) (Supplemental Table 3). Using antibodies against the germ cell lineage marker VASA (DDX4/MVH) and SFPQ, we first determined the specific localization to the nucleus in both female and male PGCs by immunofluorescence on tissue sections (Fig. 5A and B). We observed the presence of SFPQ signal on the nuclei of the VASA positive cells (PGCs), but not in the VASA negative cells (somatic cells). Thereafter, we performed IF followed by TERRA RNA-FISH to evaluate the cololocalization of SPFQ and TERRA on isolated female and male 13.5 dpc PGCs. Since the number of TERRA foci varies within female/male, we compared the percentages of colocalizing foci between male and female PGCs. Percentages were obtained from the number of TERRA-SFPQ foci divided by the total TERRA foci, and multiplied by 100). Interestingly, we observed colocalization in both female and male PGCs (Fig. 5C). However, female PGCs showed increased percentage of colocalization of TERRA-SFPQ (36%) compared to male PGCs (30%) (Fig. 5D; p=0.01), indicating although TERRA interacts with SFPQ in both sexes, there is an increased interaction in female PGCs

**Figure 5.**
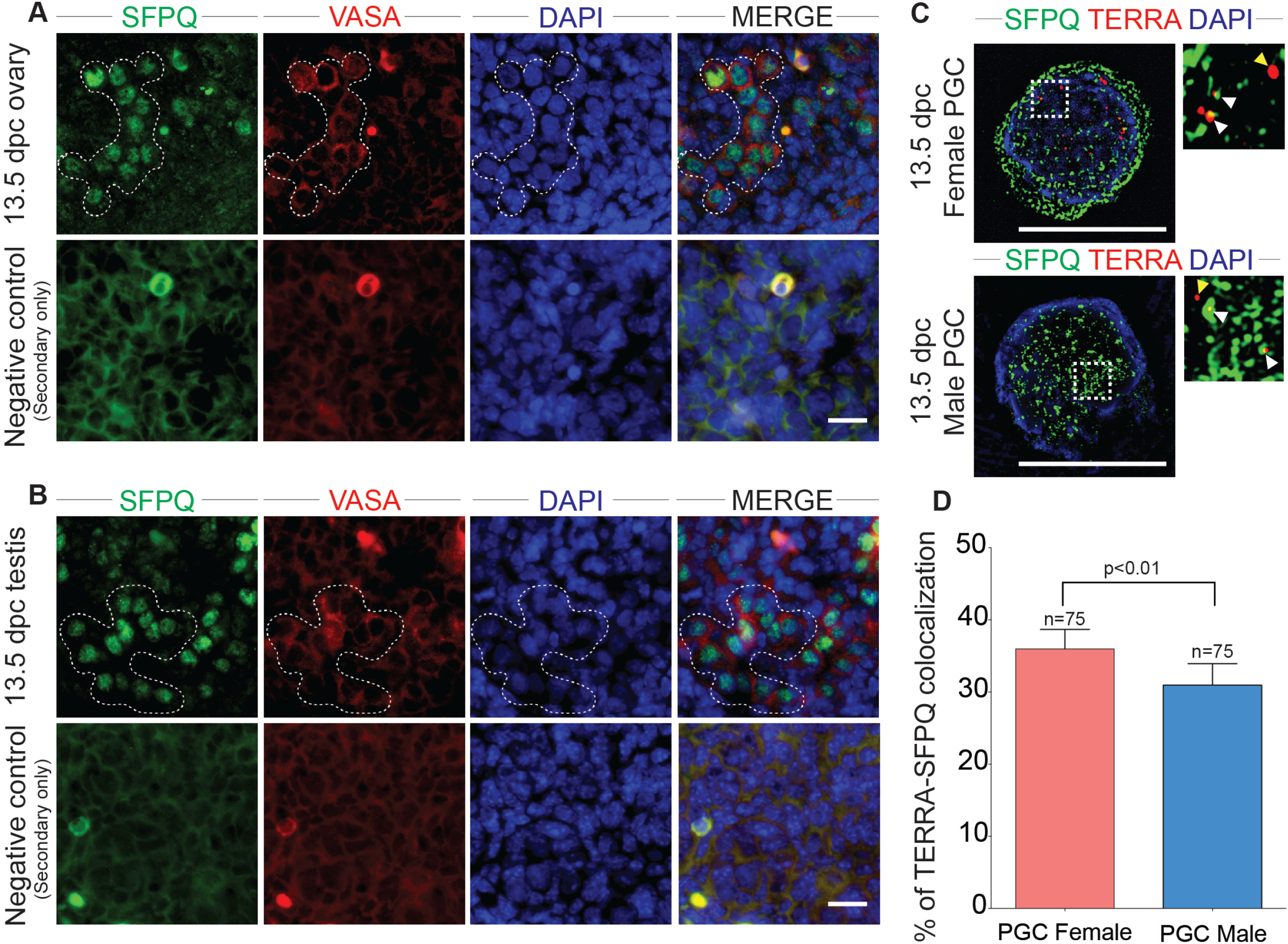
SFPQ interacts TERRA on PGCs. A) Immunofluorescence on 13.5 dpc ovary with antibodies against SFPQ (green), VASA (red) and DAPI. Scale bar 20µ; B) Immunofluorescence on 13.5 dpc testis with antibodies against SFPQ (green), VASA (red) and DAPI. Scale bar 20 µm; C) Super resolution imaging microscopy of SFPQ (green) immunofluorescence followed by TERRA RNA-FISH (red) on female and male 13.5 dpc primordial germ cells. Scale bar 10 µm; in the inset the white arrowhead shows TERRA-SFPQ interaction, yellow arrow head shows only TERRA foci D) Analysis of the percentage of colocalization of TERT-TERRA on male and female PGCs. Percentages were obtained from the number of TERRA-SFPQ foci divided by the total TERRA foci, and multiplied by 100. Statistical analysis was performed using t-test analysis. p value was set at 0.05

### NONO interacts with TERRA in somatic cells of the fetal gonad

Similar to our approach for SFPQ, above, we selected one TERRA-ineracting protein to validate on somatic cells. We analyzed the localization and interaction of TERRA with Non-POU domain-containing octamer-binding protein (NONO). First, we evaluated the localization of NONO in female and male gonads, the use of VASA antibody allowed us to identify PGCs from somatic cells. In both sexes, we observed a positive NONO staining in somatic cells with a clear absence of NONO staining on the nuclei of the VASA positive cells (Fig. 6A and 6B). Using IF followed by TERRA RNA-FISH we evaluated the colocalization of NONO with TERRA on isolated somatic cells (Fig. 5C). Colocalization analysis showed that the 42% of the TERRA foci colocalized with NONO on 13.5 female somatic cells, however on the male somatic only the 25% of the TERRA foci colocalized with NONO (percentages were obtained from the number of TERRA-NONO foci divided by the total TERRA foci, and multiplied by 100) (Fig. 5 C and D; p=0.01). These results indicate that the colocalization TERRA and NONO is restricted to the somatic cell lineages of the fetal gonad.

**Figure 6.**
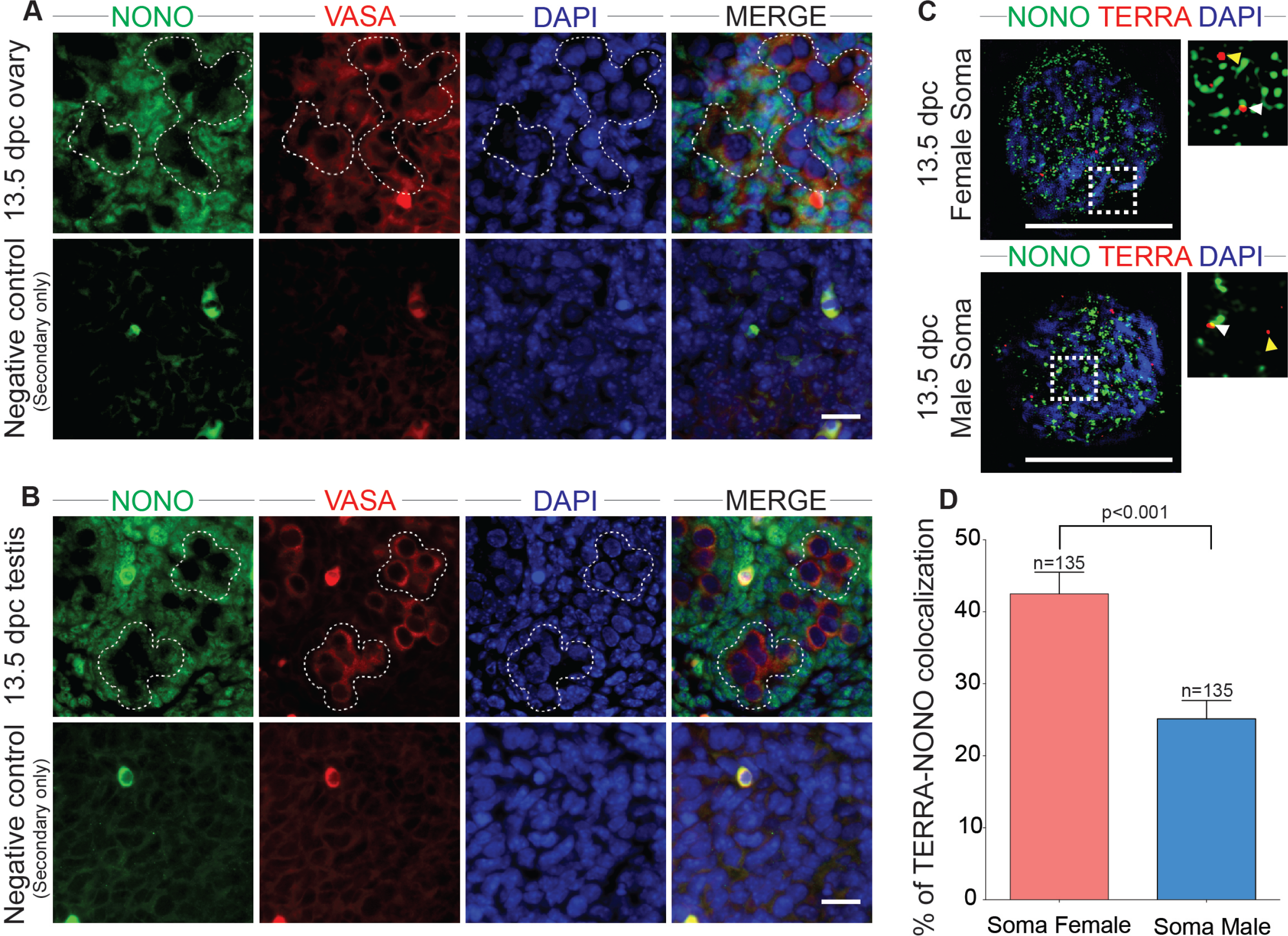
NONO interacts with TERRA on gonadal somatic cells. A) Immunofluorescence on 13.5 dpc ovary with antibodies against NONO (green), VASA (red) and DAPI. Scale bar 20 µm; B) Immunofluorescence on 13.5 dpc testis with antibodies against NONO (green), VASA (red) and DAPI. Scale bar 20 µm; C) Super resolution imaging microscopy of NONO (green) immunofluorescence followed by TERRA RNA-FISH (red) on female and male 13.5 dpc somatic cells. Scale bar 10µm; in the inset the white arrowhead shows TERRA-NONO interaction, yellow arrow head shows only TERRA foci D) Analysis of the percentage of colocalization of TERRA-NONO on male and female somatic cells. Percentages were obtained from the number of TERRA-NONO foci divided by the total TERRA foci, and multiplied by 100. Statistical analysis was performed using t-test analysis. p value was set at 0.05

## Discussion

To date, there is a very limited understanding of the function of TERRA, especially as it pertains to germ cells. Herein, we describe for first time the localization, expression and the protein interactions of TERRA in PGCs during the critical period of epigenetic reprogramming and the onset of sex determination. TERRA transcription, at least in human cells, was described to be regulated by methylation status ^15, 40^. During PGC development, genome-wide DNA demethylation in mid-gestation results in the lowest levels of methylated DNA at around 13 dpc ^21, 41^. Therefore, we hypothesized that the transcription and localization of TERRA within primordial germ cells will increase inversely to the drop in DNA methylation. At the same time, this period is associated with gonadal sex determination and the concomitant entry of female PGCs into meiosis, with male PGCs entering a period of mitotic quiescence. Thus, we suggest that this period of development would provide for key differences in TERRA regulation between the male and female germ lineages. Our results showed that both TERRA focus distribution and TERRA transcription increases with gestational age in both female and male PGCs. However, the increases are significantly higher in PGCs isolated from ovaries compared to those obtained from testis, and there is a delay by one day in the onset of TERRA transcription in males.

Sexual dimorphism in the epigenetic marker and DNA methylation patterns in PGCs has been extensively studied at different stages of gestation in the mouse ^21, 22^. In both sexes, the lowest levels of genome-wide methylation are observed at 13.5 dpc, but the recovery of the epigenetic marks takes longer in the female germ line compared to male^42^. Concommitant with the return of epigenetic marks in female PGCs is their synchronous entry into the meiotic program. Thus, the increased TERRA localization may be highly influenced by the onset of prophase I in the female germ line but we cannot discard the role of the changes in methylation status. Indeed, since one proposed function of TERRA is to regulate/suppress DNA damage repair processes, and since prophase I is characterized by the induction and orchestrated repair of hundreds of DNA double strand breaks (DSBs), it is tempting to speculate that one role of TERRA is to prevent DSB induction/repair at telomeres and instead to direct DSB events to more proximal chromosomal locations. The purpose of prophase I DSB induction/repair is to generate a highly regulated number of crossovers that serve as tethering points to maintain homologous chromosome interactions until the first meiotic division. Thus the placement of DSB events at telomeres would not be ideal for this purpose.

Interestingly, TERRA focus numbers in both male and female PGCs are significantly lower than in somatic cells, and represent a frequency that is less than a quarter of the number of telomeres present in the nucleus. In the soma, by contrast, only half the diploid number of chromosomes (40 in mouse) appear to be associated with a TERRA focus. It is important to note that somatic cells of the gonad do not undergo to epigenetic mark erasure, indicating that TERRA transcription in PGCs and somatic cells at these particular stages of development is likely to be regulated by different mechanisms. Controversial results have been published about the expression of TERRA and *Tert*, where it has been described that TERRA either recruits *Tert* to the telomere to promote its enzymatic function, or conversely TERRA is acting as a competitive inhibitor that is competing for access to telomeric DNA^17, 36^. Our results showed that in PGCs the expression of TERRA and *Tert* decreases in relation to gestational age in agreement with previous reports ^37, 38, 43^. Our results demonstrate intermittant colocalization of TERT and TERRA, and also showed that when transcription of TERRA increases the *Tert* transcription decreases. However, more studies analyzing the interaction of TERRA with TERT are required to evaluate whether TERRA affects the enzymatic activity or the transcription of *Tert* in PGCs.

A previous study had investigated TERRA-protein interactions in MEFs using a pre-existing RNA-IP technique^44^ only 41 proteins were identified as part of the TERRA interactome. Analysis of TERRA interacting proteins also has been performed using SILAC-labeled nuclear cell lysates, pooled results from 2 independent pull downs showed approximately 924 interacting proteins ^45^. More recently iDRiP was used to examine TERRA interactions in 15×10^7^ mouse embryonic stem cells and the authors report 134 interacting proteins ranging from components of the shelterin complex, to chromatin associated proteins, DNA repair proteins, and cell cycle regulators, to name just a few ^14, 18^. In all these reports, the authors use cultured somatic cells or ESCs, allowing for high cell input (≈30 million cells). By contrast, the current study analyzed TERRA interacting proteins starting with 3 million PGCs from each sex and an equivalent number of somatic cells, representing cell purifications from 300 pups for each sex. Despite the low input, we were able to identify 48 proteins were identified in female PGCs and 26 in male PGCs. 37.5 % of these proteins were shared with the iDRiP study of Chu *et al*. (2017) using ES cells ^14^. These common protein interactors included several heterogeneous ribonucleoproteins, histones, chromatin associated proteins, RNAP II and DNA repair proteins, all found in PGCs and in somatic cells. Interestingly, the SFPQ-TERRA interaction that we identify in female and male PGCs was not observed in ES cells suggesting potential specificity of this interaction for germ cells. By contrast, in somatic cells of the gonad, we identified NONO as an interactor of TERRA, and this interaction was also identified in ES Cells by Chu *et al*. (2017). The comparison between our data and previous reports suggest that there are interactions conserved in different cells types, however TERRA also have specific protein interactions depending of the cell type and at least in PGCs the sex is another determinant factor.

To validate our iDRIP data, we chose to examine further one PGC-specific (SFPQ) and somatic cell-specific (NONO) interacting protein. We did not observe clear differences in protein localization on histology sections in 13.5 dpc ovary and 13.5 dpc testis. We analyzed the colocalization of TERRA-SFPQ and TERRA-NONO, in both cases the percentage of colocalization TERRA-Protein was higher in females compared to males, indicating that at least at this specific point of development, the interaction/function of the TERRA-Protein complex is different. SFPQ is DNA- and RNA binding protein, while NONO is an RNA splicing factor ^46^. Both proteins have been implicated in a range of DNA/RNA metabolic processes, including ssDNA invasion to generate a D-loop, non-homologous end joining (NHEJ), and mRNA processing and stabilization^47^. However, their function in the developing mouse gonad including meiosis, and particularly in connection with TERRA, remains unknown.

Taken together, herein we described for first time the presence of TERRA, in mouse primordial germ cells. Our results also showed that in mouse PGCs TERRA foci number, expression and protein interactions depend on the gestational age and sex suggesting that the role of TERRA telomere dynamics is related to sexually dimorphic changes in germ cell development with age.

## Author contribution

M.A.B-E and P.E.C designed experiments. M.A.B-E, S.L.M and A.B-F carried out experiments. M.A.B-E, S.L.M, and P.E.C analyzed and interpreted data. M.A.B-E and P.E.C wrote the manuscript.

**Summary sentence:** TERRA transcription and interacting proteins during PGC development are regulated in a dynamic fashion that is dependent on gestational age and sex

## Acknowledgements

We are grateful to Dr. Jen Grenier, Dr. Rita Reig-Viader, Dr. Anand Minajigi and Dr. Jeannie Lee for their advice and protocol sharing to perform the qPCRs, RNA-FISH and iDRiP. We thank Dr. Sheng Zhang and Dr. Ruchika Bhawal from Cornell Proteomics and Mass Spectrometry Facility for his advice in development of mass spectrometry experiments. We are also thankful to Dr. Stephen Gray, Dr. Kathryn Grive, Carolyn Milano, Amanda Touey and Jeffrey Pea for providing critical feedback for the manuscript.

---

**Supplementary Table 1.** TERRA and specific subtelomeric chromosome primers

**Supplementary Table 2.** PGCs protein ratios obtained from iDRiP (m/f)

**Supplementary Table 3.** Somatic cells protein ratios obtained from iDRiP (m/f)

**SUPLEMENTARY TABLE 1.**
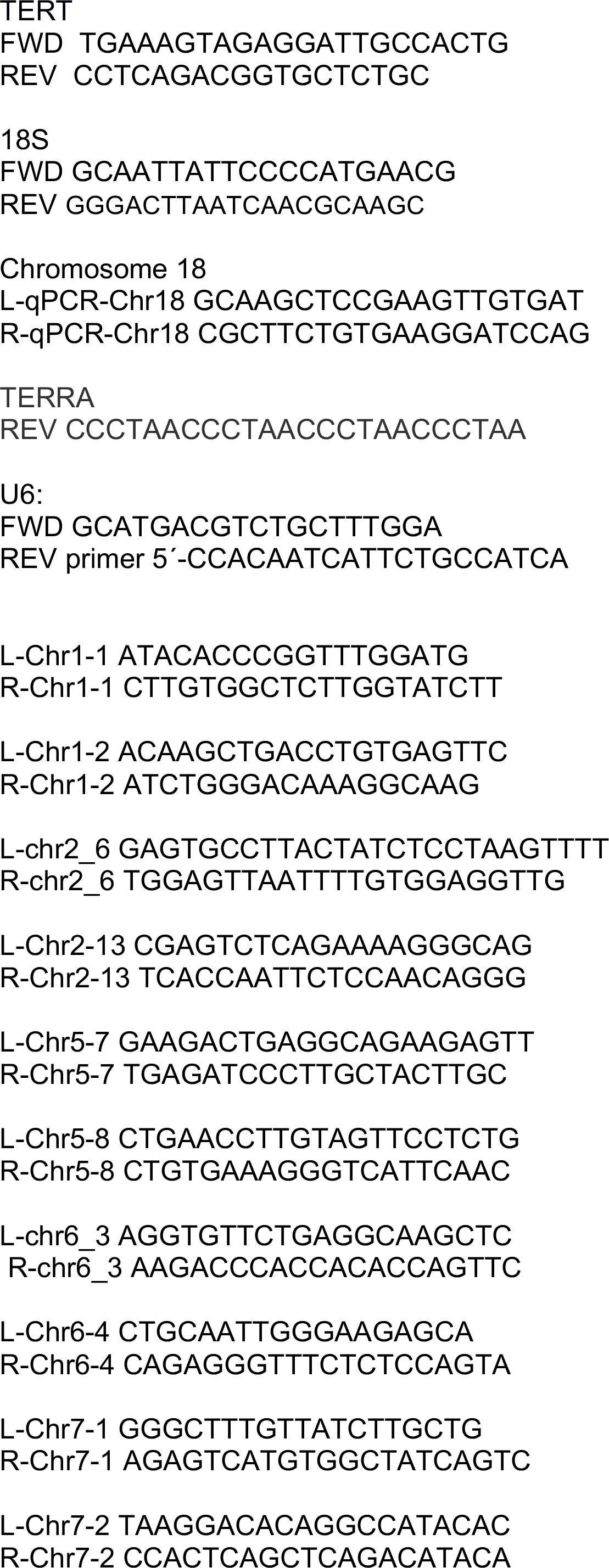
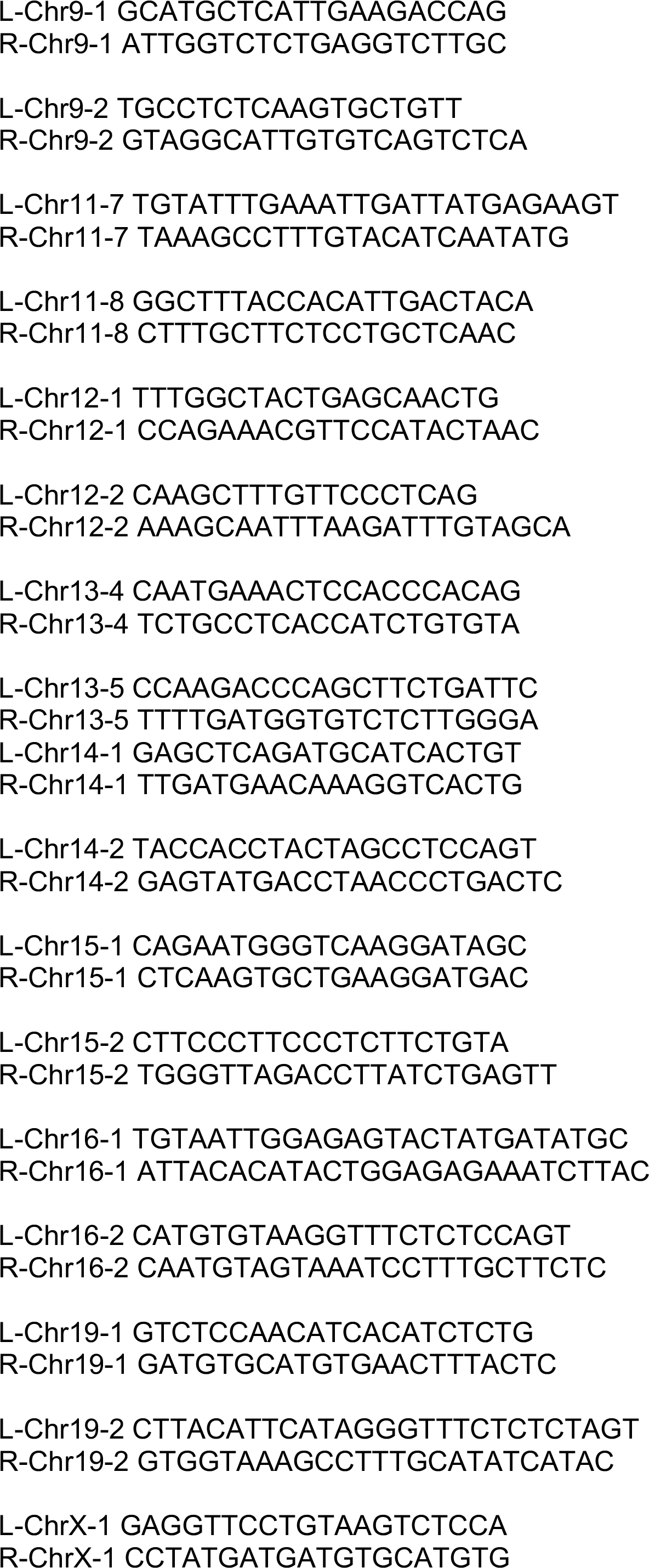
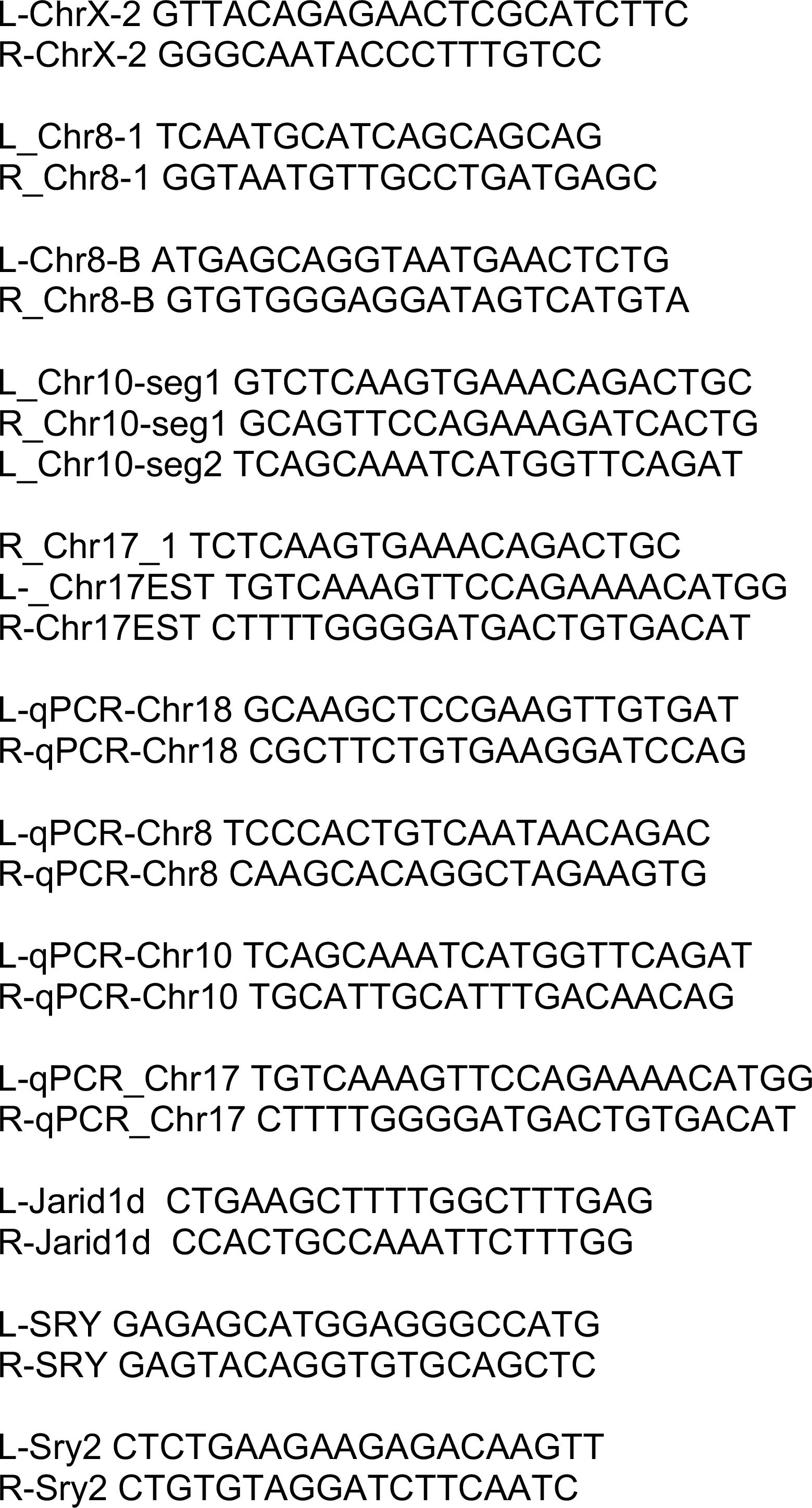
PRIMERS USED IN THIS STUDY.

**SUPPLEMENTARY TABLE 2.**
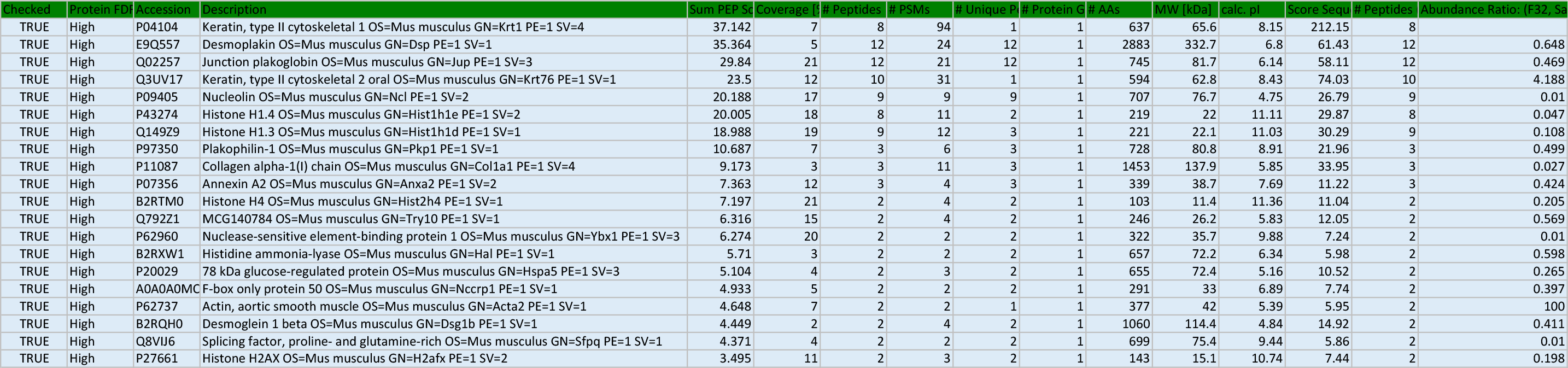
PGC PROTEIN RATIOS FROM IDRIP (m/f)

